# ArchMap: A web-based platform for reference-based analysis of single-cell datasets

**DOI:** 10.1101/2024.09.19.613883

**Authors:** Mohammad Lotfollahi, Chelsea Bright, Ronald Skorobogat, Mohammad Dehkordi, Xavier George, Simon Richter, Vladimir Shitov, Helena Melcher, Noah Nussbaumer, Annika Fomm, Yuanyu Li, Alexandra Topalova, Malte D. Luecken, Fabian J. Theis

## Abstract

Leveraging single cell reference atlases to analyse new data has brought about a paradigm shift in single cell data science akin to the first reference genome in genomics. However methods to perform this mapping require computational expertise as well as sometimes considerable compute power, and thus may exclude the researchers from this innovation who may benefit the most from it. ArchMap, a no-code query-to-reference mapping tool, removes this barrier by providing all-in-one automated mapping, cell type annotation, and collaborative features to analyse single-cell datasets from a wide range of integrated, often published, reference atlases and allows extension of atlases with the growing Human Cell Atlas and related efforts. This paves the way for a democratisation of reference mapping capabilities.

---

Large-scale integrated single-cell reference atlases have been revolutionary for understanding inter-cellular variation and achieving consensus on cell type nomenclature^1–4^. A central use of these reference models has also been to gain insights into new data using transfer learning^5–9^, similar to genome references in genomics. However, leveraging these methods requires computational know-how and, even with this know-how, setting up the complex pipelines required for atlas use is time-consuming and potentially prohibitive for users.

To address this limitation, tools that simplify the use of these methods have been developed; however these tools are either limited in their analysis capabilities or are restricted by paywalls. For example, Azimuth^7,8^ provides a graphical user interface (GUI) allowing atlas access to its users to project new single-cell data; however, the platform lacks collaborative features and mapping can only be done using methods from Seuratv4. This prohibits compatibility to atlases integrated using strictly python methods. Additionally, Fastgenomics^10^ provided reference mapping to the HLCA; however, access is no longer freely available. scGPT Hub and www.celltypist.org/ are GUIs that can also be used for reference mapping and cell type annotation, respectively; however, scGPT is a model pre-trained on diverse datasets and only fine-tuned on the user’s desired reference and query dataset and thus may introduce certain biases and remaining batch effects from other non-relevant datasets. Furthermore, celltypist on its own does not allow for removal of batch effects and thus another method or tool would need to be used to do the integration of the query into the reference.

ArchMap (Architecture Mapping; https://www.archmap.bio) is a free, no-code query-to-reference mapping framework that provides compatibility for python-based mapping methods and includes out-of-the-box cell type label transfer, uncertainty estimation, and collaborative analysis features. The fully automated approach makes query mapping accessible to users with and without coding and machine learning expertise through its intuitive GUI and access to a full code tutorial for more advanced users. A cloud-based setup, enables secure sharing of results, codeless downstream analysis, and collaborative annotation of the user’s mapping results. ArchMap also introduces metrics not currently supported in existing tools, including uncertainty quantification of label transfer that takes into account potentially correlated cell latent features and a presence score that quantifies the likelihood that a reference cell is present in a query dataset. An important use case of this metric is assessing which cell types are present or under-represented in organoids compared to an existing tissue reference. This is shown by the authors of the human neural organoid cell atlas who mapped their atlas to the atlas of the developing human brain^11^.

Archmap was developed with 4 guiding principles in mind: maximising accessibility; protecting user data privacy; ensuring utility, quality, and reproducibility; and enabling collaboration.

Traditional data analysis pipelines are prone to bottlenecks such as installation and package dependency issues, limiting usability. A cloud-based setup allows for easy access and convenience by removing the need for software installations and updates by the user and providing automatic access to CellxGene Annotate^12^ for downstream data analysis.

In addition, as users would typically leverage ArchMap during early stages of data processing, when concerns about data security and privacy are highest, ArchMap is specifically designed to keep user data protected through our google cloud implementation. All user data is encrypted during upload, download, and when stored, ensuring only authorised parties have access at any time. Furthermore, all projects deleted by the user are automatically deleted from cloud storage after 3 days.

To ensure utility of ArchMap across datasets from different tissues, Archmap is built to ingest high quality reference atlases integrated using different methods. A wide variety of atlases are available on ArchMap, including atlases accepted by the Human Cell Atlas as official tissue references^1,11,13–21^. Additionally, Scvi-hub compatibility allows for coverage of atlases with no existing preprint or publication^22^. Users who would like to use their own atlas for mapping or who would like to promote their work can utilise ArchMap’s atlas upload feature. Upon upload, the atlas is initially marked as “in revision” while it undergoes integration quality checks (Methods; Extended Data Fig. 1). If all quality checks are passed, the atlas uploader can choose whether they would like to make their atlas publicly available on the ArchMap platform or keep the atlas for private use (Fig. 1a; Atlas upload revision in Methods). Mapping to these atlases is made possible through scArches^23^, ArchMap’s backbone for query mapping. ScArches is a transfer learning framework, which can be extended to any conditional neural network-based data-integration method. Current compatible integration methods include scVI^6^, scANVI^9^, and scPoli^5^, with scVI and scANVI being consistent top performing integration methods^4^. Furthermore, to cover potential performance differences for user queries, the user has access to a variety of pre-trained classifiers for post-mapping cell type classification. These include ArchMap’s out-of-the-box XGBoost and KNN implementations, as well as scANVI and scPoli native classifiers for compatible atlases (Fig. 1b). Using 80-20 train test splits, we assessed the performance of each classifier and found KNN performs best across all atlases and cell type label granularities (Extended Data Fig. 2). To assess the overall effectiveness of the mapping, ArchMap provides mapping quality metrics. In particular, along with the Euclidean uncertainty, to quantify cell type uncertainty for uncorrelated variables, ArchMap includes Mahalanobis uncertainty, which allows for the quantification of cell type uncertainty where the latent features of the cell are correlated. These uncertainty metrics can be used to assess the reliability of the mapping and to identify novel cell states in query datasets. Additionally, a built-in presence score measures which reference cell states are present in the query dataset, which can be used to quantify the suitability of a reference for a particular query dataset (see Methods).

**Figure 1.**
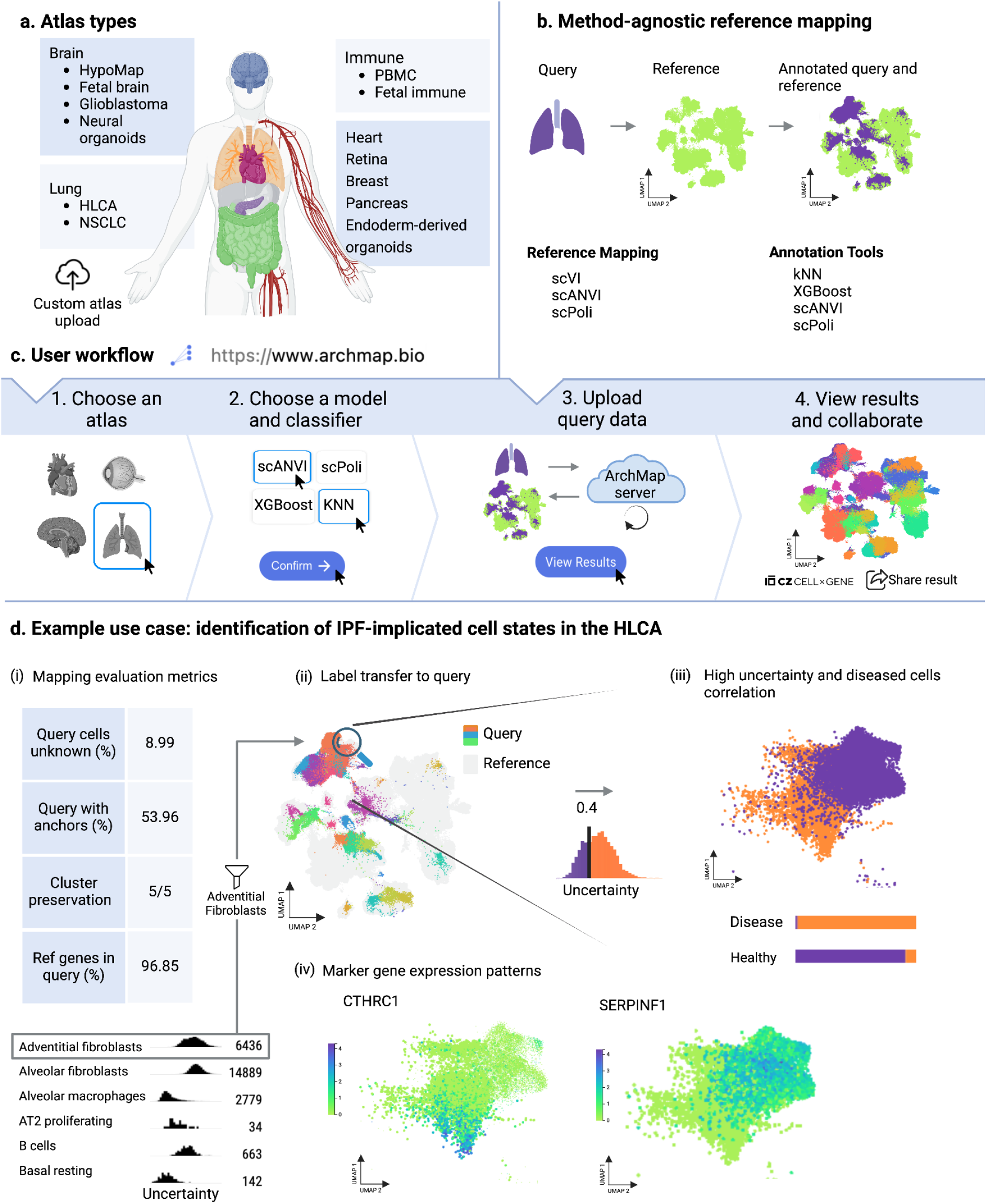
A web-based platform for automated single cell reference mapping and annotation. **a**, Atlas types available to map to on the ArchMap website. Atlas builders can upload their own atlases which are published on the website upon review. **b**, ArchMap’s backbone, the scArches query mapping method, enables reference mapping to atlases built with a variety of atlas learning methods. A selection of classifiers allow users to compare classification models and choose the best performing model for their dataset based on the percentage of query cells labelled unknown. **c**, The standard ArchMap user workflow consisting of atlas and model selection for mapping and cell type label transfer; a query upload page to drag and drop or select your desired h5ad file for mapping, as well as an outline of the prerequisites the h5ad file should satisfy before upload; and finally a results page where mappings can be visualised with a CellxGene plugin, evaluation metrics can be viewed, and your results can be downloaded. **d**, Use case mapping a single-cell dataset of human samples from patients with idiopathic pulmonary fibrosis to the HLCA. (i) Post mapping, mapping evaluation metrics can be viewed on the ArchMap website to assess the quality of the mapping. (ii) A CellxGene instance can be launched to visualise the query dataset mapped onto the reference atlas. The UMAP of the mapping is coloured by the cell type label predictions for the query cells computed by ArchMap, with the reference cells greyed out to distinguish query from reference. Cell type-specific density plots of the ArchMap-computed uncertainty scores are compared with adventitial fibroblasts shown to have a higher variability in its uncertainty scores. (iii) A correlation can be seen between the uncertainty scores of adventitial fibroblasts and the disease label of a cell. (iv) Marker gene expression patterns of CTHRC1 and SERPINF1 show higher expression in high and low uncertainty cells, respectively.

Moreover, given the interdisciplinary nature of single cell genomics, collaboration is crucial for effective research. ArchMap’s cloud infrastructure allows for easy and secure data sharing as well as collaborative annotation and data analysis with CellxGene. To balance data sharing with privacy considerations, Archmap users can choose to keep their data private or share with other users within or across teams or institutions.

## ArchMap user workflow

The standard ArchMap user workflow starts with the user choosing a fitting reference atlas and classifier for query mapping and label transfer, respectively. Upon atlas and model selection, the user is provided the necessary instructions to upload their dataset to be queried against the chosen reference model. Once the query dataset is uploaded, the user can track the processing status of each of their mapping projects in the custom “Your Mappings” page. Here, users can also view the mapping quality metrics automatically calculated by ArchMap post mapping. Once processing is completed, a CellxGene Annotate instance can be launched with the click of a button to display the joint embedding of reference and query data. In this visualisation, users can display uncertainty and presence scores calculated for each cell and make use of all CellxGene Annotate features including adding categorical cell annotations and conducting differential gene expression analysis (Fig. 1c).

### Use case: identification of IPF-implicated cell states

To demonstrate how the above workflow can be used to analyse a new dataset, we mapped a single-cell dataset of human samples from patients with idiopathic pulmonary fibrosis^24^ to the Human Lung Cell Atlas reference^1^ using ArchMap. As a first step, we see that the quality of our mapping is good as indicated by the performance metrics in Fig. 1d (Methods). The gene overlap is high at 96.85%, which is necessary to ensure a good quality mapping. The cluster preservation score is 5 out of 5, meaning that the Leiden clusters calculated on the query dataset before mapping were preserved. We see that 53.96% of query cells have a corresponding reference cell (anchor) that is considered to be in a similar biological state. Additionally, 8.99% of query cells have an uncertainty score above 0.5 and are thus labelled as “Unknown”. We note that a lower percentage of query cells with anchors and higher uncertainty scores for query cells is expected when mapping a disease dataset to a healthy reference. To continue the data analysis workflow on ArchMap, we launch Archmap’s CellxGene plugin to visualise and perform downstream analyses on the mapping results. Here, we first inspect the newly generated cell identity labels and the confidence at which these were annotated. Comparing the distribution of uncertainty scores for each cell type, we see significant variability in adventitial fibroblasts (Fig. 1d), a cell type known to be associated with IPF.^24^ Stratifying adventitial fibroblasts into low and high uncertainty groups, we highlight a correlation between high uncertainty scores and diseased cell labels (Fig. 1d). Inspecting the differential expression (DE) gene signatures associated with the uncertainty signal using ArchMap’s CellxGene plugin, we find distinct expression patterns in these two groups. Indeed, a deeper analysis of the DE signal, we find that low uncertainty fibroblasts are associated with homeostatic processes while high uncertainty fibroblasts express fibrosis signatures. For example, CTHRC1 expression, known to be unique in the fibroblasts of fibrotic lungs^24^, is elevated in high uncertainty cells (Fig. 1d). Thus ArchMap is able to detect adventitial fibroblasts as a disease cell state based on uncertainty metrics and identify driver genes such as CTHRC1 in a newly mapped IPF dataset. Importantly, this analysis can be conducted entirely through ArchMap, without the need for local data downloads or software installations.

Taken together, Archmap enables query-to-reference mapping and out-of-the-box cell type annotation for new data using existing references from a multitude of tissues. ArchMap automatically calculates various performance metrics, including uncertainty quantification to evaluate mapping quality and identify novel or diseased cells. A CellxGene plug-in allows for easy post-mapping visualization and marker gene identification. Users also have the option to share their mapping results with other users directly through ArchMap. We encourage atlas builders to promote their own work through ArchMap’s atlas upload and integration quality assessment feature. Due to server limitations, query mapping is restricted to 200 000 cells per mapping. However, users can batch their query and submit separate projects and concatenate their mapping results easily with provided Colab notebooks (see “Query requirements” in Methods). This method gives the same performance as mapping the entire query at once (Extended Data Fig. 3). We foresee ArchMap growing to accommodate multi-modal reference mapping by extending to incorporate deep learning frameworks such as MultiMIL^25^. Due to the ever increasing size of reference atlases, we also see potential in extending ArchMap to support multiple lineage atlases, which will enable users to map to a specific view/subset of an existing atlas. Furthermore, as existing atlases continue to be updated, ArchMap’s capability as an atlas repository can be further extended towards an open source atlas version control platform. As it stands, ArchMap simplifies the query-to-reference mapping process for a wide variety of users and provides a more accessible alternative to traditional mapping pipelines. With its intuitive design and collaborative, secure interface, we envision ArchMap to become a key first step for wetlab biologists to query, annotate and understand their single cell data.

## Methods

### Atlas preprocessing

Each atlas is preprocessed using the same steps to prepare it for use in the ArchMap machine learning pipeline. First, the atlas cells and genes are filtered to conform with the attributes of the respective trained model. Then, the atlas is minified^22^. In particular, the latent posterior parameters are calculated and stored in the .obsm attribute of the AnnData object. This step is done before any mapping to save computing the parameters during a query mapping, reducing compute time drastically. Additionally, the count matrix is removed from the atlas AnnData object before mapping to further reduce compute time and the launch time of the CellxGene instance. The combined reference and query count matrix is then re-added to the h5ad file containing the final joint results for download.

### Query requirements

All query requirements are outlined on the query upload page under the heading “Query Upload Requirements!!” when creating a mapping on the website. The query data file needs to fulfil certain requirements upon upload by the user. To test whether the query matches the requirements, users can make use of our tutorials for queries with either an rds or an h5ad format. The file should be of h5ad format. Furthermore, the batch/study key of the data is mandatory and should be labelled as “batch”. This is done by creating a new column in the obs data frame of the h5ad file. Users can make use of the colab notebook [here](https://colab.research.google.com/drive/11a-QpqPnBFvdB3ySQSZuoICXvuWKxQwm?usp=sharing), and referenced on the websites and in the docs to add the batch key to their h5ad file.

Furthermore, the size of the query has a limit of 200 000 cells due to limitations in server capacity. If your query exceeds this limit, you can split your query into multiple batches and create separate mapping projects. The resulting h5ad files can then be concatenated post-mapping. This procedure will not harm the mapping performance (Extended Data Fig. 3); however, it is important that all cells with the same batch label remain in the same query subset upon splitting. You can use the following linked colab notebooks to [separate](https://colab.research.google.com/drive/1-1kLHbo6sfvnOXKUbY49Nw3iCItIYky9?usp=sharing) and [concatenate](https://colab.research.google.com/drive/1voDJg4Yyz9yuubd76VaIn0fdKPsTJ6ue?usp=sharing) your query and results. These links are also referenced on the website and in the docs.

Users should also ensure that raw expression counts are saved in .X of the query AnnData object, unless otherwise specified, and that either Ensembl IDs or gene names are stored in the var_names AnnData object attribute of your query. This is dependent on the choice of atlas. Some atlases require Ensembl IDs, while others require gene names. The requirement on which label should be stored in your query is given on the query upload page of the website.

During preprocessing in the ArchMap pipeline, query genes of the uploaded query are subsetted to match the genes of the atlas and for any missing genes, expression values are padded with zeros. This may lead to inaccuracy in results based on the number of missing genes in the query. The user will be made aware of this on the website during query processing. Specifically, the user will receive a message on the frontend that communicates the percentage of shared genes between the query and reference. The user can then use this information to scrutinise the accuracy of the resulting mapping. We recommend that mappings where more than 20% of the reference genes are missing in the query should be treated with caution.

### Mapping procedure

The mapping procedure in ArchMap is outlined as follows: during the query upload, the batch key of the query is checked to make sure that it is labelled as “batch”. If not, a message is returned to the user stating that the batch key should be renamed to “batch”.

Next, the query data is preprocessed based on the model that was used for the atlas integration. For the case of scVI and scANVI, the prepare_query_anndata method for the respective model is run to add zero padding for missing genes in the query and to sort the genes to match the atlas. Subsequently, the setup_anndata method is used to map between the data fields used by the model and their locations in the query anndata. These methods prepare the query to be loaded to the model using the method load_query_data implemented in SCVI and scANVI codebase. For the case of scPoli, the preprocessing steps mentioned above for scVI and scANVI are incorporated into scPoli’s implementation of the load_query_data method.

After the query is loaded and the model is initialised with the new data, the model is trained using 100 epochs to update the reference model with the query data and obtain a joint embedding. The query latent embedding is then calculated by feeding the input query data through the updated model encoder. The query embedding is then combined with the pre-computed reference embedding and stored in the obsm anndata attribute of the joint anndata object under the key name “latent_rep”. Before combining the query with the reference, we label the query and reference cells as either “query” or “reference” in the “type” key of the obs anndata attribute to be used for visualisation in CellxGene.

### Visualisation and data download

CellxGene Annotate is used to visualise the query mapping. Users can colour their mapped query by the label transfer results saved under the category on the left hand side that ends with “prediction_knn” or “prediction_xgboost” (depending on the classifier that was chosen by the user).

Before a CellxGene instance is created for visualisation of the mapping results, the reference cells of the joint representation are subsetted to speed up computation of the UMAP and to guarantee conformity to CellxGene cell number requirements for the case of a large reference such as the PBMC atlas which has 7 million cells. The subsetting is done in a way that preserves the cell type proportions in the reference data and keeps the ratio between reference and query no smaller than 5:1. The user has the ability to download the full mapping, which is in the form of a tar.gz file consisting of the combined reference and query dataset in h5ad format and the fine-tuned model; however, the neighbour graph will need to be calculated by the user for the full data if visualisation is desired on the downloaded data. A colab notebook has been prepared [here](https://colab.research.google.com/drive/1CxyNutNUjK3SSLJbxhj_Gg89FO4Rr5EO?usp=sharing) which can be used by the user to calculate the neighbour graph of their results.

### Label transfer and uncertainty quantification

The XGBoost and KNN pre-trained classifiers were trained on each atlas and evaluated using an 80/20 split strategy. Comparing the F1 scores for the KNN, XGBoost and native (either scPoli or scANVI dependent on the model used to integrate the atlas) classifiers, KNN performed the best for all annotation levels of all atlases.

If the pre-trained classifier chosen by the user is KNN or XGBoost, the pre-trained classifier and label encoding for the user-specified atlas is downloaded from GCP. If the native scANVI or scPoli classifier is chosen, the respective instantiated model that has been updated on the query data is used for the classification.

ArchMap uses Euclidean and Mahalanobis uncertainty to quantify the label transfer uncertainty. The Euclidean uncertainty calculated for a cell is based on the Euclidean distance between that cell and its 15 nearest neighbours. This score is used to help the user identify novel cell states in the query data. We have added a separate column, named “{cell_type_key}_prediction_{classifer_name}_filtered_by_uncert>0.5”, in the output that labels cells as “unknown” above a certain uncertainty threshold. Here, cell_type_key is the key in the obs attribute of the cell type level mapped to and classifer_name is the name of the chosen classifier (e.g. knn). The threshold is set to 0.5 and is based on the way the Euclidean uncertainty is calculated. The Euclidean uncertainty of the query cells is calculated using the weights of a weighted k-nearest neighbour classifier trained on the reference embedding. The KNN classifier is applied to the query embedding, where the distances to the 15 nearest reference cells are measured for each query cell. A Gaussian weighting function is used to assign more importance to nearer reference cells, and is given by

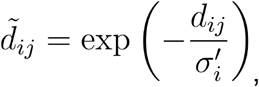

where *s*_*ij*_ is the Euclidean distance between the *i*-th query cell and the *j*-th nearest reference cell and 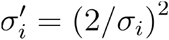 where *σ*_*i*_ is the standard deviation of the distances to the 15 nearest reference cells.

The weight *w*_*ij*_ corresponding to the *j*-th nearest reference cell of the *i*-th query cell is normalized by the sum over all *K* neighbors so that the weights for each query cell sum to 1:

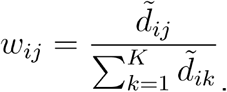

For each unique cell type label from the reference nearest neighbours, a probability for that label is calculated by summing the corresponding reference cell weights. The reference label with the highest probability is then used as the label for the query cell. To calculate the uncertainty for that query cell we take 1 − *p*, where *p* is the probability of the reference label used for label transfer. Thus, we set a threshold of 0.5 since a probability below 0.5 is more uncertain than certain and no label stands out as a strong prediction.

For the case that users are interested in considering novel cells in the query dataset, we have also included in the output a column with all transferred labels for each query cell without any threshold used. We named this column “{cell_type_key}_prediction_{classifer_name}”. This allows users to also obtain transferred labels for query cells with uncertainty greater than 0.5. Furthermore the “{cell_type_key}_uncertainty_euclidean” column can be used to view the exact uncertainty scores for all query cells.

The Mahalanobis uncertainty is based on the Mahalanobis distance between a query cell and the means of the Gaussian components from a Gaussian mixture model of the reference embedding, where the number of components is equal to the number of cell types. In particular, the Mahalanobis uncertainty is given by

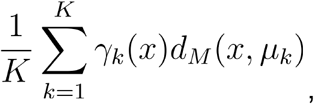

where 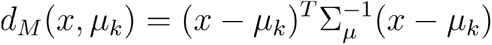 is the Mahalanobis distance between a query cell and the mean of the *k*-th Gaussian component. Here, 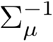 is the inverse covariance matrix derived from the means of the Gaussian components. Furthermore, *γ*_*k*_(*x*) is the probability of query cell *x* belonging to component *k*. This probability is the posterior probability calculated using Bayes’ Theorem:

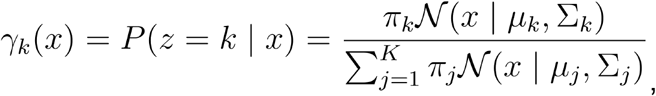

where 𝒩 (*x* | µ_*k*_, Σ_*k*_) is the probability density function of the *k*-th multivariate Gaussian distribution and *π*_*k*_ is the prior probability of Gaussian component *k*.

Mahalanobis uncertainty accounts for potential correlation between the latent features of the cells. This is done by rescaling the latent features such that the covariance between any two correlated latent features is removed. If there are no correlations between the features, the distance *d*_*M*_ is equal to the Euclidean distance.

We note that both the Mahalanobis uncertainty and Euclidean uncertainty seem to be able to effectively identify potential novel or disease cell states for the case of the IPF dataset used to map to the HLCA in this paper. However, we have chosen to allow for both uncertainty scores to be calculated to allow for users to compare the uncertainty scores for their specific query datasets, making the tool more robust to embeddings that may still retain correlations between features.

### Performance metrics

During processing, performance metrics to quantify the quality of the mapping are calculated. These metrics include the percentage of reference genes in the query data, the percentage of cells labelled “Unknown”, the percentage of query cells with anchors, and the cluster preservation score. For the percentage of reference genes in the query data, a value less than 85% may contribute to poor mapping quality. The percentage of cells that are labelled as “Unknown” during cell type label transfer is based on the Euclidean uncertainty score for each query cell. A query cell is classified as having an “Unknown” cell type if its Euclidean uncertainty score is above 0.5. A value of 0.5 is used here to take into account query datasets where cells of the same type may look different from the reference atlas (e.g. when the query data consists of cells from diseased tissue.

The percentage of query cells with anchors represents query cells that have at least one mutual nearest neighbour among the cells of the reference dataset. These mutual nearest neighbours are termed anchors. A lower percentage of query cells with anchors may not necessarily mean poor mapping quality. A lower score may be returned for query datasets from the same tissue as the reference, but with a large proportion of cells from diseased donors. For example, mapping a dataset consisting of samples from healthy tissue^26^ to the HLCA gives a percentage of query cells with anchors of 63%. However, mapping a dataset consisting of samples from diseased tissue^24^ to the HLCA gives a percentage of query cells with anchors of 54%. We also note that the percentage of anchor cells is affected by the number of cells in the query and reference data. Thus this metric should not be used to compare mapping quality across query or reference datasets, but only across models and classifiers. The purpose of this metric is to convey the success of the mapping and the suitability of the chosen atlas as a reference for the specific query data.

The cluster preservation score assesses how well Leiden clustering of the query dataset is preserved after the mapping process. The mean entropy of cluster labels within each neighbourhood is computed for both the original and integrated query. The median of the differences in mean entropy between the original and integrated query is then calculated. Scores are scaled between 0 and 5 with 5 being the best.

### Presence score

The presence score is based on the weighted shared nearest neighbours between the reference and query cells. A high presence score for a reference cell indicates a high frequency and likelihood of such a cell state or one similar being in the query. This complements the uncertainty scores given to the query cells.

### Query mapping onto HLCA

ArchMap can be used for the identification and validation of disease-specific cell states. To show this we mapped a single-cell dataset of human samples from healthy donors and patients with idiopathic pulmonary fibrosis to the Human Lung Cell Atlas reference and used the resulting uncertainty scores calculated in the ArchMap server to identify potential disease-associated cell states and analyse marker genes.

The percentage of reference genes in the query data is 96.85%. The percentage of cells labelled “Unknown” is 8.99% and the percentage of query cells with anchors is 53.96%. The lower percentage of anchors may be a result of the query data containing a large portion of cells from diseased donors, while the reference only contains cells from healthy donors. The cluster preservation score is 5 out of 5, implying perfect preservation of clusters in the query.

Once the user’s query data has been mapped onto the reference using the ArchMap server, the mapping can be visualized in CellxGene. The embedding can be coloured based on query vs. reference and based on the cell type label transfer predictions. To quantify the uncertainty of the cell type label transfer, the Mahalanobis uncertainty is used. Colouring by this uncertainty metric, the user can identify clusters of interest that warrant further analysis. Adventitial fibroblasts can be seen as one of the cell type clusters with the most varied uncertainty scores. We expect that clusters with both low and high uncertainty contain both healthy and diseased cells.

Upon filtering, we split the cluster of interest into two groups: (1) all reference cells and all query cells with uncertainty scores less than 0.4 and (2) all query cells with uncertainty scores greater or equal to 0.4. Investigating the correlation between the uncertainty scores and the ground truth cell labels of healthy and diseased, we find a positive correlation between higher uncertainty scores and cells from diseased donors. This suggests that the uncertainty scores can be used as a starting point to identify disease-associated cell states (Fig. 1d).

Differential expression analysis can be performed in CellxGene to compare low and high uncertainty groups. In Fig. 1d, the embedding of adventitial fibroblasts are coloured by the gene expression of CTHRC1 and SERPINF1, two genes from the top 40 most differentially expressed genes for the high and low uncertainty group, respectively.

### Access to scvi-hub atlases

On ArchMap, by default, users have access to the latest versions of all scANVI- and scVI-integrated scvi-hub atlases (upload of scPoli-integrated atlases is currently not available on scvi-hub). For the case where users would like to use an earlier atlas version on scvi-hub, we have included a [tutorial](https://archmap-docu.readthedocs.io/en/latest/tutorials/scvi_hub.html) in the ArchMap docs that walks the user through downloading different versions of an atlas from scvi-hub. The user can then upload this atlas to ArchMap to do their mapping on their desired atlas version.

Publicly available scvi-hub atlases that are compatible with scArches mapping include all compartments of the multiple-organ Tabula Sapiens atlas and the Human Lung Cell Atlas. These atlases are published and thus can be considered as trustworthy. However, for atlases uploaded to scvi-hub in the future, we will include only atlases with a pre-print, publication, or model evaluation results in the model card in scvi-hub, as described [here](https://docs.scvi-tools.org/en/1.0.0/tutorials/notebooks/scvi_hub_upload_and_large_files.html#model-evaluation). For users who would like to map their data to an scvi-hub atlas that does not meet these requirements can upload the desired atlas and run ArchMap’s benchmarking process as described [here](https://archmap-docu.readthedocs.io/en/latest/beta_feature/index.html#atlas-revision-stage).

### Atlas upload revision

To support community uploads of reference atlases, a platform user must first be an admin. In order to become an admin, the user must reach out to ArchMap’s development team. This can be done by checking the box “Request for Beta feature” upon registration to ArchMap. Once approved, the user can start contributing by providing additional atlases. A user-uploaded atlas must first go through a revision process before it is publicly available on the website. The revision process consists of the following evaluation steps: (1) benchmarking the user uploaded model used for integration with scVI, scPoli with prototype learning, scPoli without prototype learning, and PCA (no batch effect removal). Benchmarking is done using scIB^4^ (2) Comparing the KNN classifier F1 scores between classifiers trained on the latent representations taken from the user integrated model, scVI, scPoli with prototype learning, scPoli without prototype learning and PCA. After uploading an atlas, a user is able to evaluate the quality of their atlas on the ArchMap website (see documentation for full details of the benchmarking process). Once the atlas has been successfully benchmarked, the user will be able download the benchmarking report from the website and can send this report to the ArchMap team at. This report contains only the benchmarking evaluation as described in points (1) and (2) above. Thus, no private data is shared through email. Once the report has been successfully reviewed to ensure integration quality, the atlas uploader can choose whether they want to make their atlas available to the public for mapping or to keep their atlas private.

### Framework architecture

Archmap’s backend is written in Express.js, running on a node.js server. The machine learning pipeline is a stand-alone service, primarily based on the scArches package for reference mapping. The pipeline is written in Python.

The backend as well as the machine learning pipeline are hosted on GCP (Google Cloud Platform). The backend is primarily running on Google App Engine. Google Cloud Storage is used to store persistent data such as project mapping results and annotation files. The machine learning pipeline as well as ephemeral CellxGene visualisation instances are run on Google Cloud Run.

### Running ArchMap locally

We have added detailed information on how a user can install ArchMap locally on our [GitHub](https://github.com/theislab/archmap?tab=readme-ov-file#install). For users who are only interested in the mapping pipeline or who are not familiar with Node.js, we have also provided a [tutorial](https://archmap-docu.readthedocs.io/en/latest/tutorials/map_local.html) in the form of a Python notebook. This tutorial provides the user with the exact mapping pipeline that is used in ArchMap, but can be run locally. The user only needs to create an environment by running a provided bash script and then running the notebook. By default, the tutorial is set to map a provided query to the HLCA. If the user would like to use their own data and a different reference, they can download the desired reference from ArchMap and change the input and output paths in the tutorial notebook.

### Number of epochs for mapping

For all query-to-reference mappings, a maximum of 100 epochs are used for fine-tuning the reference model. This number was chosen by running mappings of both healthy and disease query data, with approximately 70 000 and 118 000 cells respectively, to the HLCA and increasing the number of epochs from 1 to 1000. We evaluated the mappings using the reconstruction loss, scIB metrics4 and UMAP embeddings (Extended Data Fig. 4). We found that reconstruction loss for both query datasets plateaus after approximately 100 epochs (Extended Data Fig. 4a). Furthermore, for the healthy query, we see all scIB metrics continue to increase up until approximately 250 epochs, at which point the kBET and PCR comparison decrease slightly. After this decrease at approximately 500 epochs they then remain constant, along with all other metrics. For the disease query we see that past 100 epochs, we begin to see a decrease in PCR comparison, kBET, and KMeans ARI, potentially attributed to over-integration of the query. Thus, we have chosen a maximum of 100 epochs since for the majority of scIB metrics, we see no further improvement past this number (Extended Data Fig. 4b). Furthermore, for both the reconstruction loss and the UMAP embeddings, we see no change or improvement in integration when training for more than 100 epochs (Extended Data Fig. 4c).

### Number of overlapping genes between query and reference

During mapping, only genes used to build the reference atlas model can be used to map a query dataset to this atlas, requiring the genes of query datasets to be subsetted to match the atlas gene set. To better understand the effect of this subsetting on query to reference mapping, we benchmarked query mapping quality under varying overlap of profiled genes in the HLCA reference (see Supplementary Note 1). This experiment has allowed us to provide a more informative guideline to users mapping their query to a reference atlas on ArchMap. Based on our benchmark, we recommend that the overlap of genes between the query and reference should be no lower than 80%.

## Code availability

The machine learning pipeline for mapping query data to reference atlases and implementing all downstream computation tasks is available at theislab/archmap_dataatprocessing_minify_scanvi(github.com)).

The code for the frontend and backend implementation of ArchMap is available at theislab/archmap(github.com)).

The ArchMap website is available at https://www.archmap.bio.

## Author contributions

M.L. conceived the study with contributions from F.J.T. M.L., M.D.L, and F.J.T. supervised the project. M.L., R.S., C.B., X.R., S.R., M.D., V.S., and A.T. designed and implemented ArchMap and all supporting documentation. H.M., N.N., A.F., and Y.L. conducted the benchmarking experiments in the extended data figures. C.B. wrote the manuscript, conducted the ArchMap use case, and finalized the project with guidance, contributions, and approval from M.D.L., F.J.T., and M.L.

## Acknowledgements

The authors thank Anastasia Litinetskaya for valuable feedback on the project and paper. We thank Anastasia Litinetskaya, Leander Dony, and Raphael Kfuri Rubens for beta testing the website. We thank Sergei Rybakov for lending his expertise on the scArches framework. This work was funded by the Chan Zuckerberg Initiative (Grant ID 230826). V.A.S. is supported by the Helmholtz Association under the joint research school “Munich School for Data Science - MUDS”

## Competing interests

M.L. consults for Santa Anna Bio owns interests in Relation Therapeutics and is a scientific cofounder and part-time employee at AIVIVO. F.J.T. consults for Immunai Inc., Singularity Bio B.V., CytoReason Ltd. and has ownership interest in Dermagnostix GmbH and Cellarity. The remaining authors declare no competing interests.

## Extended data

**Extended Data Fig. 1 ArchMap’s atlas upload evaluation pipeline results for the HLCA**

**a**, Comparison of KNN classifier F1 scores between classifiers trained on the latent representations taken from the user integrated model (scANVI), scVI, scPoli without prototype loss, scPoli with prototype loss, and PCA. Cell type labels used for ground truth were “ann_level_5” (finest annotation level) **b**, Overview of integration methods used to integrate HLCA ranked by overall score. **c**, Scatter plot comparing batch correction vs. bio conservation scores for each method. Red dashed lines represent the scores for PCA.

**Extended Data Fig. 2 Comparing F1 scores of KNN, XGBoost, and native cell type label classifiers for pancreas, HLCA, and HNOCA**

Comparison of F1 scores between KNN, XGBoost, and scANVI native classifiers for the **a**, HLCA using “ann_level_3” as ground truth and **b**, pancreas using “cell_type”. For both atlases, KNN gives the best performance. KNN also gives the best performance for finer cell type labels. **c**, Comparison of F1 scores between KNN, XGBoost, and scPoli native classifiers for HEOCA using “level_2” cell type labels as ground truth. Again KNN also gives the best performance for finer cell type labels.

**Extended Data Fig. 3 Comparing mapping and label transfer performance of separately mapped query batches vs. a single query mapping containing all batches**

Comparing label transfer performance of separately mapped query batches vs. a single query mapping containing all batches for the **a**, HLCA and **b**, HNOCA. Both methods give the same results for cell type label transfer. **c**, Comparing scIB metrics between the two scenarios mapped to the HLCA.

**Extended Data Fig. 4 Benchmarking number of epochs used for reference mapping**

Benchmark of the number of epochs to be used for query-to-reference mapping using both disease and healthy queries mapped to the HLCA. Mapping quality is evaluated using the **a**, reconstruction loss **b**, scIB metrics using the scib-metrics package, and **c**, UMAP embeddings.

## Notes

### Summary of Updates

We have updated the figure as there was an error in the way that the figure was coloured. We have also updated the extended figures and some text based on updated features of the tool presented.

